# Similar gut bacterial microbiota in two fruit-feeding moth pests collected from different host species and locations

**DOI:** 10.1101/2020.04.06.028886

**Authors:** Qiang Gong, Li-Jun Cao, Jin-Cui Chen, Ya-Jun Gong, De-Qiang Pu, Qiong Huang, Ary Anthony Hoffmann, Shu-Jun Wei

## Abstract

Numerous gut microbes are associated with insects, but their composition remains largely unknown for many insect groups, along with factors influencing their composition. Here, we compared gut bacterial microbiota of two co-occurring agricultural pests, the peach fruit moth (PFM) and the oriental fruit moth (OFM), collected from different orchards and host plant species. Gut microbiota of both species was mainly composed of bacteria from Proteobacteria, followed by Firmicutes. The two species shared bacteria from the genera *Pseudomonas*, *Gluconobacter*, *Acetobacter*, and *Pantoea*, although endosymbiotic *Wolbachia* was the most abundant genus in PFM and *Lactobacillus* was the most abundant in OFM. PFM tended to have lower diversity and richness of gut microbiota than OFM; however, only some of the comparisons were statistically significant. Orchards can influence gut microbiota in terms of richness, particularly for PFM, but not so much for diversity parameters. Functional prediction of gut microbiota showed that the top pathways are amino acid metabolism, translation, and membrane transport in both species, but their abundance varied between the two moth species. These results show that two fruit moths share many features of gut microbiota, and the bacterial species are relatively stable within moth species even when they use different host plants. Our study suggests that fruit-feeding behavior may play a role in shaping gut microbiota of the two fruit moths, which may provide microbial targets for pest control.

**Importance:** Understanding the associated microbes with insects can point to new targets for pest control. Here we compared bacterial community in the gut of two co-occurring agricultural pests, the peach fruit moth (PFM) and the oriental fruit moth (OFM), collected from different orchards and host plant species. We found that the bacterial genera *Pseudomonas*, *Gluconobacter*, *Acetobacter*, and *Pantoea* are abundant and shared in two moths. The composition of the bacterial species is relatively stable within moth species even when they use different host plants, indicating that the gut microbiota community in the PFM and OFM is likely to be related to their fruit-feeding behavior. The findings have implications for developing novel pest control approaches by targeting gut microbes associated with the two moths.

## Introduction

Many microorganisms have become adapted to their insect hosts, forming close mutualistic relationships (1, 2). These microbes play important roles for their hosts, such as in the digestion and nutrient absorption of host food, protection against pathogens, and enhancement of immunity (3–5). The study of insect microorganisms can point to new approaches for the control of agricultural pests and human disease vectors as well as increasing the value of economically important insects, particularly by modifying the symbiotic relationship between symbionts and their hosts (6, 7).

The community of microorganisms living in insects can be affected by many environmental factors (1, 8, 9). In particular, gut bacteria of different insects can vary greatly in number, composition, distribution, and function for species adapted to different hosts and living in different habitats (10). Moreover, there can be a dynamic interaction between bacteria living in the gut and the environment as indicated by the acquisition and loss of *Serratia* symbiotica strains in aphids (11)

Moths include some of the most damaging agricultural and forest pests from the order Lepidoptera. Being holometabolous, moths are characterized by different life stages and can vary in their gut microbiota during development (12–14). Many moths are polyphagous, having a wide range of diets, which represent one of the factors impacting bacterial communities in this group (13, 15). However, while moths represent useful model species to understand the determinants of gut microbiota across life stages (16), there is limited information on variation in their microbiota.

Here we focus on the peach fruit moth (PFM), *Carposina sasakii*, and the oriental fruit moth (OFM), *Grapholita molesta*, common agricultural moth pests damaging many economically important fruit crops, such as apple, pear, and peach (17–20). Larvae of both these species bore into and feed on fruit, while OFM can also bore into tree shoots prior to pupation. These species usually co-occur in the same orchard and sometimes on the same fruit (21–23). The concealed lifestyle and wide range of host plant species used by these species make them useful to understand factors affecting their gut microbiota. Previous studies found that larvae of these two moths harbor a high diversity and richness of bacteria (24, 25), but it is not yet clear the two species are more likely to share the same gut microbores when they live in the same orchard and on the same host plant species.

We examined gut bacterial microbiota in co-occurring PFM and OFM collected from the same host plant species, with the microbiota characterized using the V3-V4 variable region of the 16S rRNA gene. We aimed to examine the relative contribution of moth species, host plant, and other factors related to variation among orchards to microbial composition.

## Results

### Community composition of the gut microbiota in PFM and OFM

The average number of sequencing reads for each sample was 4927 after filtering (**Table S1**). Rarefaction curves from both the original sequencing data sets and randomly subsampled data sets showed that the curves of all samples tended to be flat, indicating that the amount of sequencing data is enough to reflect most of the microbial diversity information in the samples (**Fig. S1**). In total, 294 OTU were clustered, attributed to 13 phyla and 176 genera and 234 species for both hosts, among which 234 OTUs belonging to 203 species and 284 OTUs belonging to 228 species were identified for PFM and OFM respectively (**Table S2**).

At the phylum level, OTUs in both species were mainly attributed to Proteobacteria (98.4% in PFM, 89.2% in OFM), followed by Firmicutes (1.06% in PFM, 8.87% in OFM) (**Fig. 1a**). At the genus level, OTUs of PFM were mainly annotated to *Wolbachia* (62.06%), *Pseudomonas* (19.09%), *Gluconobacter* (6.98%), *Acetobacter* (4.05%), and *Pantoea* (3.59%), while OTUs of OFM were mainly annotated to *Pseudomonas* (49.96%), *Gluconobacter* (12.53%), *Pantoea* (10.70%), *Lactobacillus* (7.65%), *Acetobacter* (6.61%) (**Fig. 1b** and **Fig. S2**). Similar patterns were found at the species level where these could be identified (**Fig. 1c, Table S3**).

**Fig. 1.**
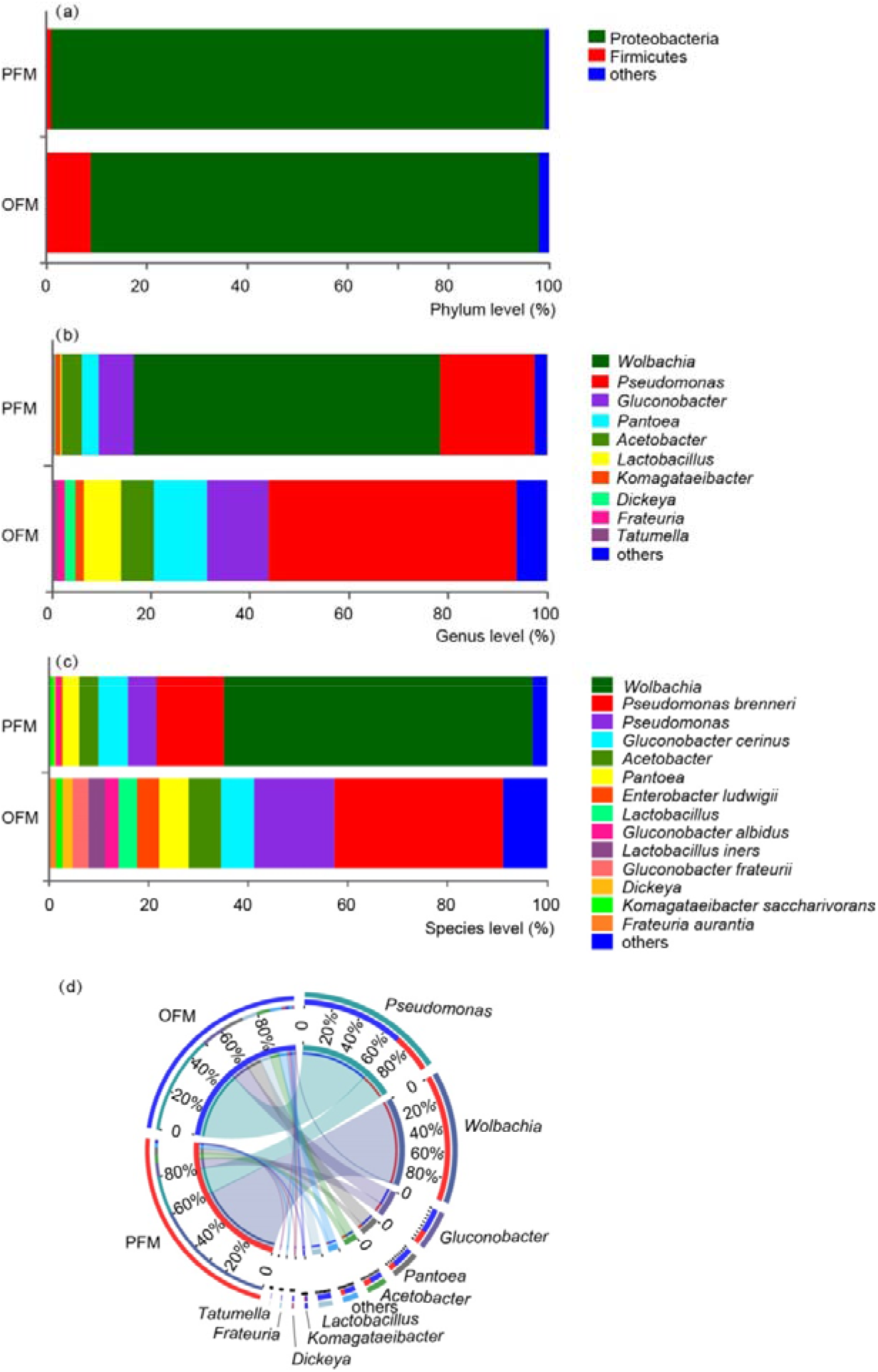
Microbial composition identified in the peach fruit moth (PFM) *Carposina sasakii* and the oriental fruit moth (OFM) *Grapholita molesta*. Community composition of the microbiome on phylum (a), genus (b), and species (c) levels for the OFM and PFM. (d) The cooccurrence relation graph describes the abundance of correspondence between samples and species. Each unit was represented by one color.

The core bacterial community at the genus level for each species was identified by comparing individuals from different orchards (**Table S4**). For PFM, 24 core genera were identified from the four orchards sampled (**Fig. 2a**), the most common of which were *Wolbachia* (67.07%), followed by *Pseudomonas* (20.63%), *Gluconobacter* (7.54%) and *Pantoea* (3.88%) (**Fig. 2b**); for OFM, 33 core genera were identified from five orchards (**Fig. 2c**), the most common of which were *Pseudomonas* (59.96%), followed by *Gluconobacter* (15.03%), *Pantoea* (12.84%), and *Lactobacillus* (9.18%) (**Fig. 2d**).

**Fig. 2.**
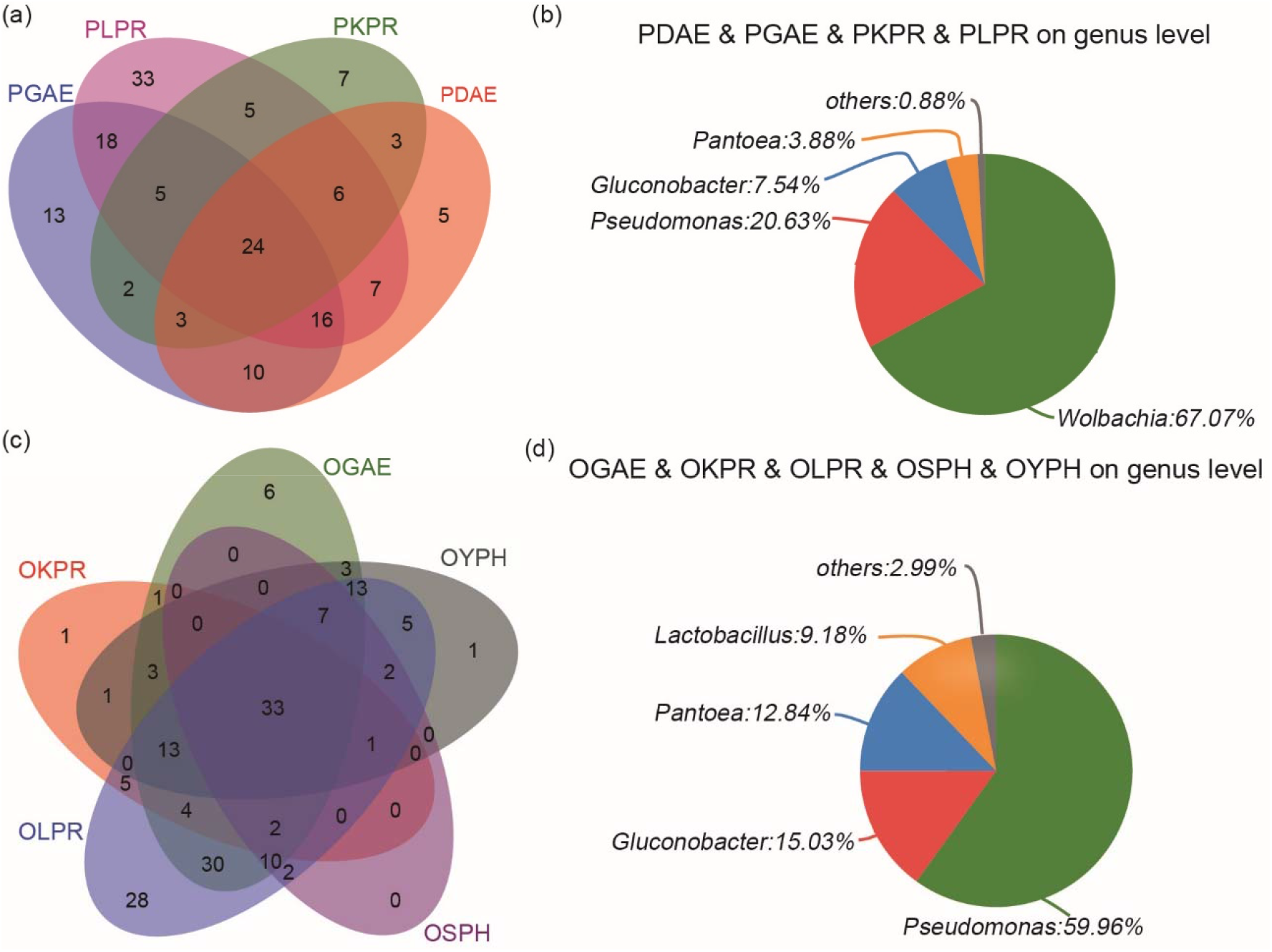
Core bacteria of the same species from different hosts and different orchards. (a) Venn diagram at the genus level of PFM in four orchards. (b) Composition of 24 core genera found in all four orchard samples. (c) Venn diagram at the genus level of OFM from 5 orchards. (d) Composition of 33 core genera found in all five orchard samples. See Table 1 for the codes.

In summary, Proteobacteria was the most abundant phylum for both host species. There were many common bacteria from the genus *Pseudomonas*, *Gluconobacter*, *Acetobacter* and *Pantoea*, although in PFM *Wolbachia* was the most abundant genus followed by *Pseudomonas*. *Pseudomonas* was the most abundant genus in OFM, and *Lactobacillus* and Dickeya were abundant in OFM but not in found in PFM (**Figs. 1d, 2b, 2d**).

### Comparison on gut microbiota between PFM and OFM

When gut bacterial microbiota was compared between all samples of PFM and OFM, in terms of alpha diversity, there was no significant difference in OTU richness between PFM and OFM (P_ace_= 0.57, P_chao_ = 0.121, P_sobs_ = 0.014) (**Fig. 3a-b, Tables S5**, and **S6**) but significantly lower diversity in PFM than in OFM (P_shannon_ = 0.002, P_simpson_ = 0.004) (**Fig. 3c-d, Tables S5** and **S6**). In terms of beta diversity, PFM and OFM individuals divided into two groups in the PCoA analysis, although outlier samples were identified (**Fig. 3e**, **Table S7**).

**Fig. 3.**
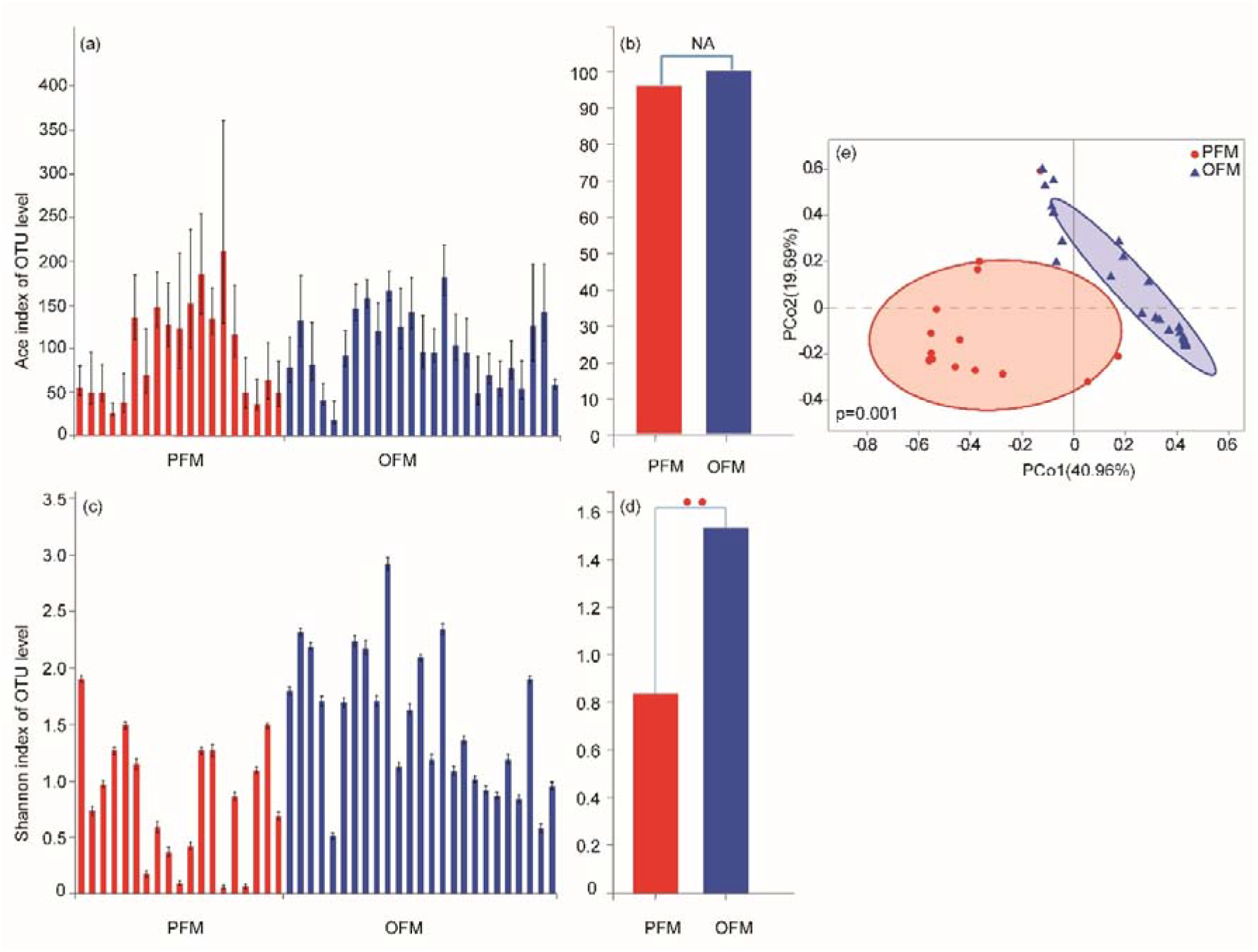
Comparison of gut bacterial microbiota between the peach fruit moth (PFM) *Carposina sasakii* and the oriental fruit moth (OFM) *Grapholita molesta* individuals for alpha and beta diversity. (a and c) Community richness and diversity by Ace and Shannon index for the OTU level between the two species. (b and d) Wilcoxon rank-sum test of the difference between OFM and PFM individuals for ACE and Shannon indices (*p* > 0.05 is marked as NA, 0.01 < *p* ≤ 0.05 is marked as *, 0.001 < *p* ≤ 0.01 is marked as **, and *p* ≤ 0.001 is marked as ***). (e) Beta diversity of the microbiome between two species estimated by PCoA analysis at the genus level. PCo1 and PCo2 are the first two principle components, while the values on the x- and y-axis are proportions explained by corresponding components, respectively (PERMANOVA test with 999 permutations, *p* = 0.001, see Table S5 for values).

**Fig. 4.**
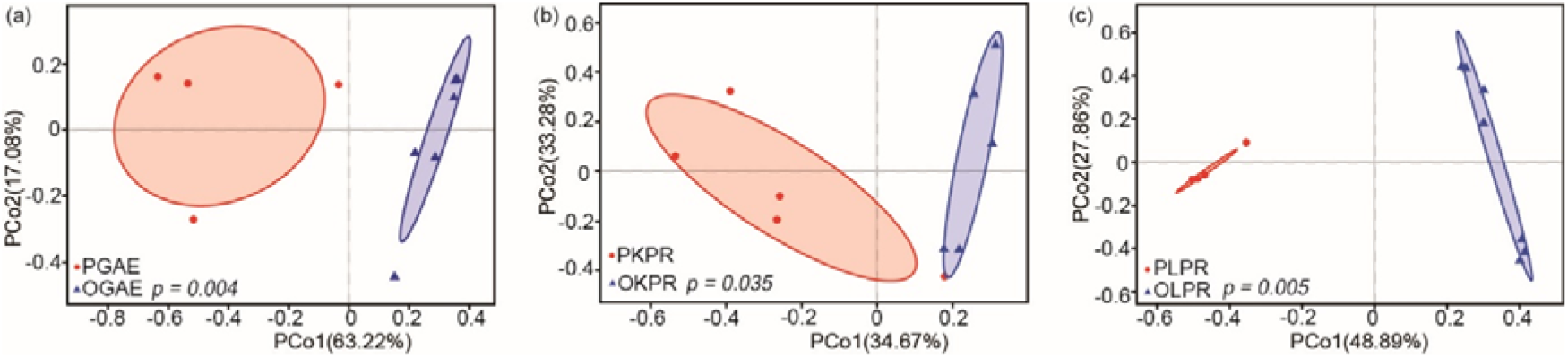
Beta diversity of gut bacterial microbiota between the peach fruit moth (PFM) *Carposina sasakii* and the oriental fruit moth (OFM) *Grapholita molesta* from the same host plant and orchard. PERMANOVA was performed to determine the differences among groups. Sampled were collected from apple orchard estimated by PCoA analysis on the genus level. (b) Sampled were collected from pear orchard. (c) Sampled were collected from another orchard of pear (PERMANOVA test with 999 permutations, see Table S5 for values).

We then compared gut bacterial microbiota between three pairs of PFM and OFM populations collected from the same host species and the same orchard. In terms of alpha diversity, OFM usually had higher richness and diversity except for one paired richness comparison collected from apple (**Fig. S3a, Table S5**). For pear, patterns were consistent, but only one of the three comparisons was statistically different in diversity **(Fig. S3h**). In terms of beta diversity, individuals of PFM and OFM from the same habitat could be clustered into different groups in the PCoA analysis (**Fig. S3c, f, i**), with individuals collected from pear showing the clearest separation (**Fig. S3i, Table S7**).

### Influence of orchard on gut microbiota within species

First, we compared the gut microbiota of the same insect species collected from different host plant species and different orchards to examine the effect of orchard but relaxing the host plant species. PFM individuals from four orchards and OFM individuals from five orchards were analyzed. For PFM, four of six pairs of orchard comparisons had significant differences in richness, while one of the six pairs showed difference in diversity (**Fig. S4**). For OFM, two of 10 pairs of orchard comparisons were significantly different for richness, but none were significant for diversity (**Fig. S5**). For overall comparison, there was no significant difference in any measure of richness or diversity in either species (χ^2^ = 18/18/18/14.36/18, df = 18/18/18/14/18, *p* > 0.4231 for Ace, Shannon, Simpson, Sobs and Chao in PFM; χ^2^ = 24, df = 24/24/24/23/24, *p* > 0.4038 for Ace, Shannon, Simpson, Sobs and Chao in OFM). Second, we compared the gut microbiota of the same species and host plant from different orchards to test the effect of the orchard by fixing the host plant. Two pairs of PFMs from pear and apple and two pairs of OFM from pear and peach shoot hosts were used for analysis. In terms of alpha diversity (Ace), a significant difference in richness was found in both paired PFM comparisons (**Figs. S6a, S6g**) and one of the two OFM comparisons (**Fig. S6d**), while a significant difference in Shannon’s index was found in one of the two PFM comparisons (**Fig. S6b**) but not in the OFM comparisons (**Figs. S6h, S6k**). In terms of beta diversity, individuals from different orchards of the same species were not clearly separated in the PCoA analyses (**Fig. 5, Figs. S6c, f, i, l**).

**Fig. 5.**
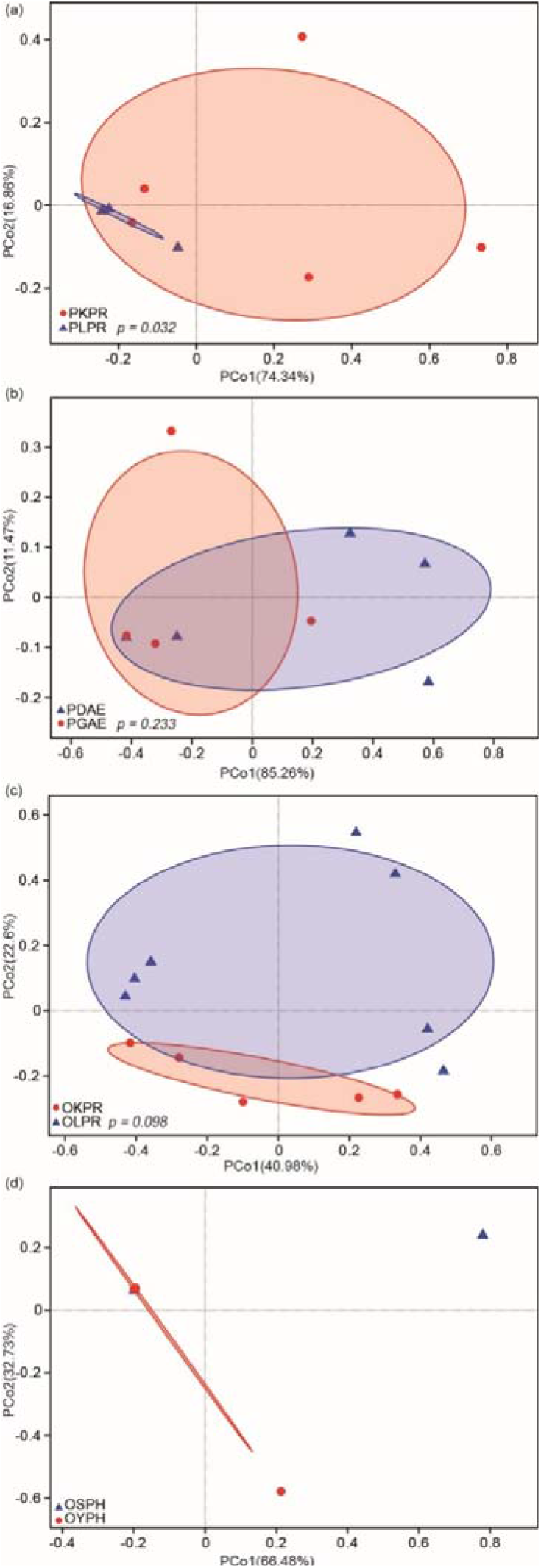
Effects of environmental differences on beta diversity of gut microbiota in the peach fruit moth (PFM) *Carposina sasakii* and the oriental fruit moth (OFM) *Grapholita molesta* between different orchards in the same insect species and the same host species (PERMANOVA test with 999 permutations, Table S6).

While these results suggest that orchard can affect the composition of gut microbiota in PFM and OFM, the effect is relatively small, particularly as shown in the beta diversity analysis. Orchard had a higher impact on richness than on diversity, and PFM tended to vary more among orchards with the same host than OFM.

### Function prediction of gut microbiota

At level 1, functions of the gut microbiota were mainly annotated to pathways of metabolism, genetic information processing, environmental information processing, and cellular processing. At level 2, the top pathways were amino acid metabolism, translation and membrane transport (**Table 2**). It can be seen in the COG (Clusters of Orthologous Groups) function annotation that the functions of gut microbiota of OFM and PFM were annotated to the same pathway, but the abundance of the same pathway was different (**Fig. 6a**). Among the top 10 functions in KEGG (Kyoto Encyclopedia of Genes and Genomes) annotations pathway level 3, there was a significant difference between OFM and PFM (*p* < 0.0001). ABC transporters and two-component system were significantly higher in OFM than PFM, and the remaining eight were higher in PFM than OFM (**Fig. 6b**).

**Table 1.**
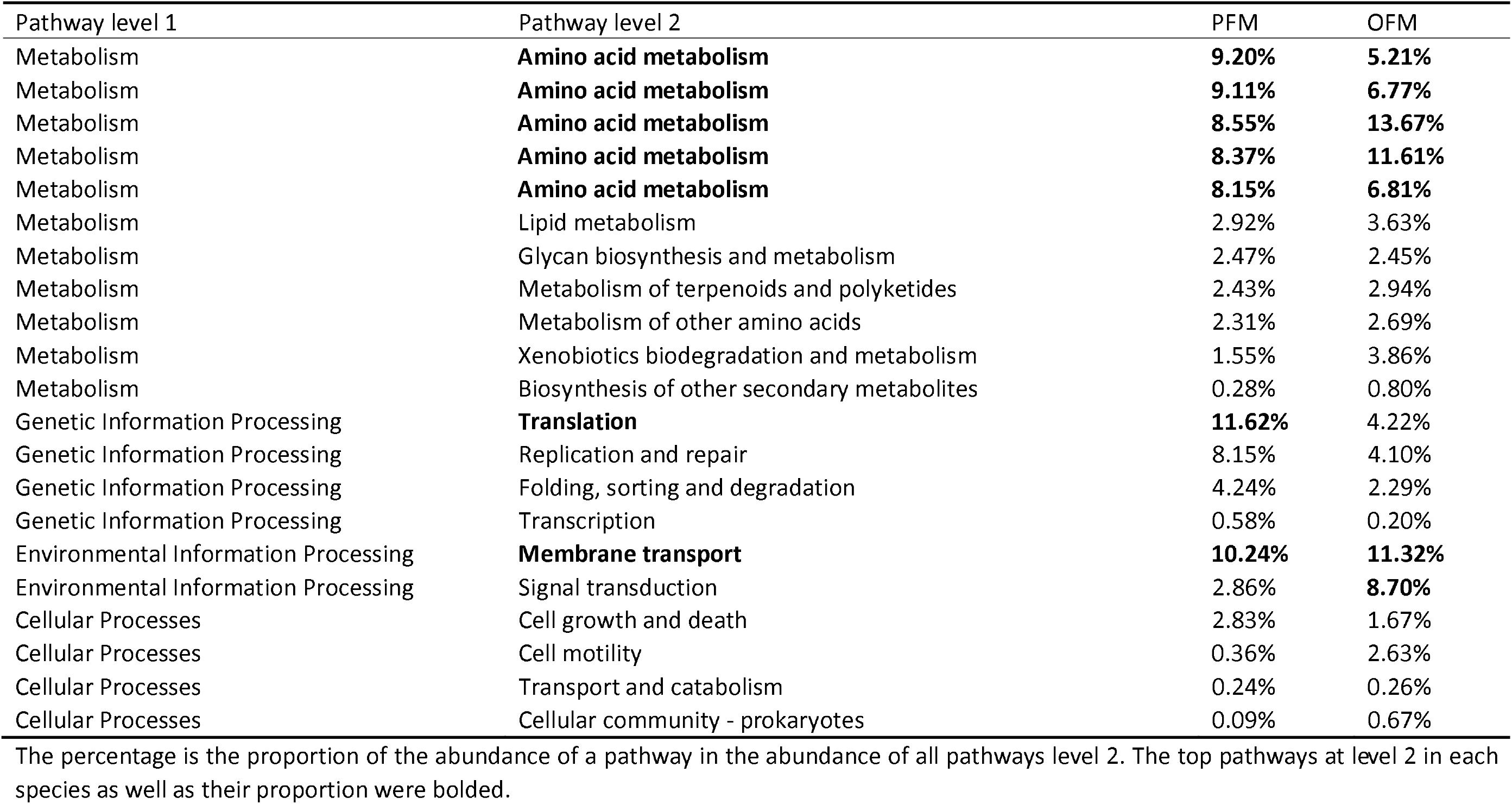
Enrichment of KEGG pathways for gut bacterial microbiota of the peach fruit moth (PFM) *Carposina sasakii* and the oriental fruit moth FM) *Grapholita molesta*.

**Table 2.** Samples of the peach fruit moth (PFM) *Carposina sasakii* and the oriental fruit moth (OFM) *Grapholita molesta* used in the study

| Code | Species | Collecting location | Host | Coordinate | NO. |
| --- | --- | --- | --- | --- | --- |
| PKPR | PFM | Kaosanji of Pinggu district (K) | Pear (PR) | 40°12′N, 117°19′E | 5 |
| OKPR | OFM |  |  |  | 5 |
| PLPR | PFM | Lvfulong of Yanqing district (L) | Pear (PR) | 40°32′N, 116°4′E | 5 |
| OLPR | OFM |  |  |  | 7 |
| PGAE | PFM | Liugou of Yanqing district (G) | Apple (AE) | 40°27′N, 116°6′E | 4 |
| OGAE | OFM |  |  |  | 6 |
| PDAE | PFM | Dafengying of Yanqing district (D) | Apple (AE) | 40°26′N, 115°54′E | 5 |
| OYPH | OFM | Linguosuo of Haidian district (Y) | Peach shoot (PH) | 39°58′N, 116°13′E | 5 |
| OSPH | OFM | Shuangxin of Haidian district (S) | Peach shoot (PH) | 39°57′N, 116°12′E | 2 |

All samples were collected from the Beijing area, China. NO., the number of individuals used for 16S rRNA gene sequencing.

**Fig. 6.**
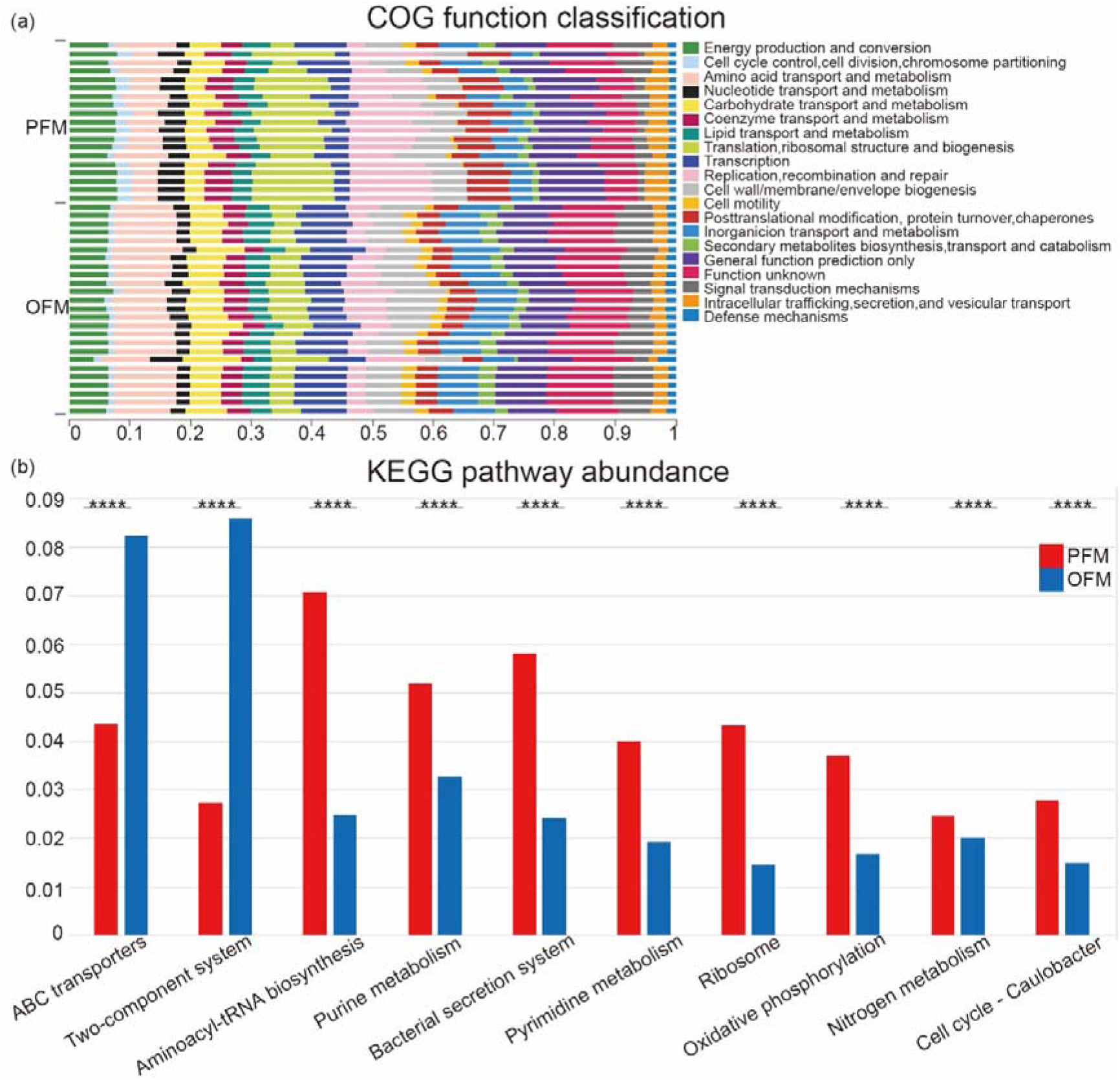
Functional predictions of gut microbiota in the peach fruit moth (PFM) *Carposina sasakii* and the oriental fruit moth (OFM) *Grapholita molesta*. (a) COG function classification; KEGG pathway abundance of top 10 KEGG pathways at level 3 and the statistical difference between PFM and OFM (*p* ≤ 0.0001 is marked as ****).

## Discussion

### Comparison of gut microbiota from two fruit borers

In this study, we found that the gut microbiota of PFM and OFM was dominated by Proteobacteria and Firmicutes, which is similar to the situation found in Y. Liu et al. (25) and Y. Li et al. (24), and in other lepidopterans such as *Lymantria dispar*, *Helicoverpa armigera*, and *Bombyx mori* (26–29). However, there was a difference between PFM and OFM and other lepidopterans at the genus level. OTUs from both PFM and OFM was dominated by *Pseudomonas*, *Gluconobacter*, *Acetobacter*, and *Pantoea*. In contrast, in silkworms, *Aureimonas*, *Methylobacterium*, *Rhizobium*, *Sphingomonas*, *Propionibacterium*, *Pseudomonas*, and *Microbacterium* were the most common genera (29). The results suggest that PFM and OFM gut microbes had a similar composition, but they are different from those of the *Bombyx mori*, which has a different diet. Our results support the notion that dietary adaptation has led to different intestinal microorganisms and symbiotic interactions (30), although more moth species with different hosts (fruit, leaf tissue, and so on) need to be included in such comparisons.

We also found some differences between the two species examined here, where OTUs of PFM were dominated by *Wolbachia*, and OTUs of OFM were dominated by *Lactobacillus*. When we focused on the gut microbes of PFM and OFM from the same host and the same orchard, this pattern was also found: *Wolbachia* was unique to PFM, while *Lactobacillus* was abundant in OFM and rare in PFM. Perhaps this difference in species might generate phenotypic differences among the species for traits such as pesticide resistance. For instance, insecticide-treated resistant strains of the diamondback moth *Plutella xylostella* had more *Lactobacillales* and the less common taxa *Pseudomonadales* and *Xanthomonadales* as well as fewer *Enterobacteriales* compared with a susceptible strain (31). The OFM microbiota might contribute to resistance, although living in fruit they would be less affected by pesticides than *Plutella xylostella* larvae feeding on leaves. The comparison of microbes of PFM and OFM in three orchards showed that there was no large difference in microbial richness and diversity between PFM and OFM, but the PCA analysis highlighted differences in species composition, with host type clearly being a major determinant of gut microorganisms.

### Influence of orchard and host species on gut microbiota

Microbial communities can vary among host locations, both in terms of community diversity and community structure (32). In our study, there were differences in microbial richness in larvae from the same species collected from different orchards with the same type of fruit (PLPR/PKPR, PDAE/PGAE, OLPR/OKPR, Table S5), which suggests an impact of orchard habitat on microbial richness. Differences in microbial diversity have also been noted in studies on other insects, such as in comparisons of *Drosophila* between indoor and wild environments (33). However, the gut microorganisms in neither PFM nor OFM could be clearly separated by orchard or fruit type in the PCoA analysis, suggesting that host species rather than location plays a more important role in microbial community composition.

### Wolbachia in PFM

*Wolbachia* is an intracellular endosymbiont rather than a gut bacterium, but it can be found in the gut wall of species (34). The role of *Wolbachia* in PFM is unclear; it is common in Lepidoptera (35) where its effects have mostly not been characterized in species although in Lepidoptera it can cause a variety of effects on host reproduction including cytoplasmic incompatibility, feminization and male-killing (36–38) and increases the susceptibility of its host to baculovirus (39). These effects have not yet been investigated in PFM and require a comparison of *Wolbachia* infected and uninfected individuals for fitness as well as crosses to establish reproductive effects.

Of particular interest from the perspective of the current study is whether *Wolbachia* might influence the gut microbiota. *Wolbachia* may lead to decreased microbial diversity due to competitive behavior (40), which may contribute to the lower diversity of gut microbiota in PFM than that of OFM. In *Drosophila melanogaster*, *Wolbachia* can reduce the richness of *Acetobacter* (41), but this group was not at a low abundance in PFM. Whether *Wolbachia* in PFM influences, other microbiota requires a comparison of *Wolbachia* infected and *Wolbachia* free lines, which might be generated through antibiotic treatment or by taking advantage of natural polymorphism in infection status within natural populations (42).

### Implications for pest management

The insect-associated microbes provide new targets for developing novel pest control methods (6, 16, 43, 44). The first step to find the potential bacterial targets is to investigate the bacterial community, its impact on the pests, and its stability. We found that the community of the gut microbiota were relatively stable within moth species in spite of host fruit differences for microbes such as *Pseudomonas*, *Pantoea*, *Lactobacillus*, *Gluconobacter*, and *Acetobacter*. Functional analysis showed that three of the ten most abundant functions were environmental signaling processes, and others involve metabolism, genetic information processing, and cellular processes. These functional classes suggest that gut bacteria have a clear interaction with host processes in the intestinal environment. Among the abundant bacteria taxa, *Pseudomonas brenneri* plays a prominent role in the removal of heavy metals (45). This species is significantly more abundant in OFM than PFM and may contribute to the different ratios of ABC transporters and the Two-component system. *Gluconobacter cerinus* was another species present in PFM and OFM, which may have a beneficial role as in the case of fruit flies where it can affect reproduction (46). *Pantoea* is a highly diverse genus that can cause plant diseases and human diseases but also have functions in habitat restoration and pesticide degradation (47). Functional studies of these bacteria may help to identify potential targets for developing control methods of these two fruit moths.

We also note that the two fruit moths share many gut bacterial taxa. The similar composition of gut bacterial microbiota indicates functions related to the common biology of both species, particularly in terms of the fruit-feeding larvae. These larvae bore into fruit or shoots soon after egg hatching, reducing their likelihood of exposure to environmental bacteria when compared to the leaf-feeding moths. In the fruit-feeding spotted wing drosophila, *Drosophila suzukii*, the gut microbiota provides nutrition by providing protein for their hosts (48). Larvae of fruit moths often feed on immature fruits, which are rich in compounds such as organic acids and tannins. The tannins are endogenous inhibitors of the growth of numerous species of pests by negatively effecting the metabolism of insects (49). We found that the most abundant function of the gut microbiota in both species were metabolic processes. There are examples of gut microbiota in lepidopteran hosts helping to detoxify host toxins (50, 51), but whether the fruit moths need microbes to help them to detoxify defensive chemicals is unclear. Nevertheless, the gut microbiota community in the PFM and OFM is likely to be related to their fruit-feeding behavior, and further tests of such hypotheses may provide insights into the development of novel control approaches.

## Materials and methods

### Sample collection and DNA extraction

We sampled three pairs of PFM and OFM populations from the same host plant and orchard, as well as one PFM population from another apple orchard, and two OFM populations from two peach orchards infesting tree shoots (**Table 2**). We collected potentially infested pears and apples and peach shoots from the field and kept them in the conditioned laboratory under 25 ± 1 °C, 60% ± 5% humidity, and a photoperiod of 16 h light: 8 h dark. Fifth instar larvae were collected when they came out from the collected hosts. Species were identified by morphology and kept in a clean 1.5 ml tube for 24 hours to clean out the feces by starvation. Then, larvae were frozen in liquid nitrogen and stored in a −80 °C refrigerator prior to usage. We examined the gut microbiota of 19 PFM and 25 OFM individuals (**Table 2**).

Prior to DNA extraction, larvae were washed three times, with 75% alcohol, and then washed three times with sterile water. The whole gut tissue was dissected and homogenized in a 1.5 ml tube by grinding manually. Total DNA was extracted from single samples using the E.Z.N.A.^®^ Bacterial DNA Kit (Omega Bio-tek, GA, U.S.) according to manufacturer’s protocol. The concentration and quality of the extracted DNA were determined by a NanoDrop 2000 UV-vis spectrophotometer (Thermo Scientific, Wilmington, USA) and gel electrophoresis on 1% agarose.

### 16S rRNA gene amplification and sequencing

We used the V3-V4 hypervariable regions of the bacterial 16S ribosomal RNA (rRNA) gene to examine the gut microbiota of PFM and OFM. A 468-bp target gene segment was amplified by primer pair of 338F (5’-ACTCCTACGGGAGGCAGCAG-3’) and 806R (5’-GGACTACHVGGGTWTCTAAT-3’) (52). For PCR reaction, 20 μL of the mixture was prepared, including 5 x FastPfu reaction buffer, 250 μM dNTPs 1 U FastPfu Polymerase (Transgene, Beijing, China), 200 nM of each prime (Majorbio, Shanghai, China), 1 μL of template DNA and DNA-free water. The PCR reaction involved a single denaturation step at 95 °C for 3 min, followed by 27 cycles of 95 °C for 30 s, 55 °C for 30 s, 72 °C for 45 s, and finished after a final extension at 72 °C for 10 min. The PCR products were run on a 2% (w/v) agarose gel and those with correct size were excised and purified with a AxyPrep DNA gel extraction kit (Axygen Biosciences, Union City, CA, USA). Illumina Miseq sequencing libraries were constructed using the TruSeqTM DNA Sample Prep Kit (San Diego, CA, USA) for the purified 16S PCR products, and sequenced on an Illumina MiSeq (San Diego, CA, USA) to obtain 300-bp paired-end reads.

### Quality control and OTU identification

Raw data from Illumina MiSeq sequencing were demultiplexed to obtain sequencing data for each sample. The quality of raw data was checked by FASTQC version 0.19.6 (53); low-quality data were trimmed and filtered by Trimmomatic version 0.36 (54). Paired-end reads were merged by FLASH version 1.2.11 (55) to generate unpaired longer reads with the following criteria: (i) the reads were truncated at any site receiving an average quality score < 20 over a 50 bp sliding window; (ii) primers were exactly matched allowing two nucleotide mismatching, and reads containing ambiguous bases were removed; (iii) only paired-end reads whose overlap longer than 10 bp were merged.

Operational taxonomic units (OTUs) were clustered with a 97% similarity threshold using UPARSE version 7.0.1090 (56), and chimeric sequences were identified and removed using UCHIME algorithm in USEARCH version 7.0 (57). The taxonomy of each 16S rRNA gene sequence was analyzed by a naïve Bayesian classifier of Ribosomal Database Project version 2.11 (58) against the SILVA rRNA database (59). To avoid the influence of sequencing depth in samples, sequences from difference samples were rarefied to the same depth. Sample sequence extraction and species screening of OTU were conducted in accordance with the following conditions: (i) removal of mitochondrial and chloroplast sequences; (ii) retention of only OTUs with sequence depth greater than or equal to five in at least three samples in subsequent analyses.

### Diversity analysis

For alpha diversity, community richness indexes (sobs, chao, and ace) and community diversity indexes (Shannon, Simpson, and Pd) were estimated. The software Mothur (60) was used to calculate the alpha diversity index under different random sampling, and the *ggplot2* R package was used to draw the rarefaction curves. The Wilcoxon rank-sum test was used to compare statistical differences between different groups, while the Kruskal-Wallis rank sum test was used for overall comparison to examine the species, host, and orchard effects. In the beta diversity analysis, principal coordinates analysis (PCoA) was conducted based on a Bray-Curtis dissimilarity matrix computed from the samples. For group comparisons, a non-parametric multivariate statistical test, permutational multivariate analysis of variance (PERMANOVA), was conducted based on the Bray-Curtis dissimilarity matrix, using Qiime and the R package *vegan* (61).

### Functional analysis

We used PICRUSt version 1.1.4 (62) to predict the function of the gut microbiota from PFM and OFM. The OTU abundance table was first normalized by removing the effect of the 16S rRNA gene copy numbers (GCNs). The COG (Clusters of Orthologous Groups) family information was obtained according to the Greengene id version gg_13_5 (63) corresponding to each OTU. The description information of each COG and its function information was parsed based on the eggNOG (evolutionary genealogy of genes: Non-supervised Orthologous Groups) database v5.0 (64). The 16S rRNA taxonomic lineage based on the SILVA rRNA database (59) was transformed into the taxonomic lineage of prokaryotes in the KEGG (Kyoto Encyclopedia of Genes and Genomes) database Release 92.0 (65) through Tax4Fun (66), and the 16S rRNA gene sequence was functionally annotated. A Wilcoxon rank-sum test was used to compare the statistical difference between OFM and PFM for the 10 most abundant pathways at level 3.

## Acknowledgements

Funding for this study was provided jointly by the National Key Research and Development Program of China (2019YFD1002102), the Beijing Key Laboratory of Environmentally Friendly Pest Management on Northern Fruits (BZ0432) and BAAFS-UOM Joint Laboratory on Pest Control Research. SJW conceived and designed research. DQP, YJG, JCC and QH collected the samples. QG and LJC conducted experiments. QG, SJW and LJC analyzed data. QG, SJW and AAH wrote the manuscript and discussed the results.

