## Supplementary material for "Similar gut bacterial microbiota in two fruit-feeding moth pests collected from different host species and locations": Tables S1-S7 and Figures S1-S6

**Table S1** Basic statistics on 16S rRNA gene sequencing of the gut bacterial microbiota for the peach fruit moth (PFM) *Carposina sasakii* and the oriental fruit moth (OFM) *Grapholita molesta* using Illumina Miseq platform. * two primers used for sequencing.

| Item | Data |
|---|---|
| Amplified region | 338F-806R* |
| Samples number | 44 |
| Passed QC (bp) | 905462037 |
| Average length (bp) | 418.2527697 |
| Total raw reads | 2164868 |
| Average reads | 49201 |
| Minimum read count | 33359 |
| Maximum read count | 71332 |
| Total chloroplast reads | 206585 |
| Total mitochondrial reads | 14499 |
| Average reads (chloroplast & mitochondria sequence removed) | 5025 |
| Minimum read count (chloroplast & mitochondrial sequence removed) | 0 |
| Maximum read count (chloroplast & mitochondrial sequence removed) | 65066 |
| Subsample read count (after filtering) | 4927 |

**Table S2** Statistics on number of OTUs and their classification identified for the peach fruit moth (PFM) *Carposina sasakii* and the oriental fruit moth (OFM) *Grapholita molesta*.

| Data type | Group | Sample | Phylum | Class | Order | Family | Genus | Species | OTU |
|---|---|---|---|---|---|---|---|---|---|
| Raw data | ALL | 44 | 39 | 84 | 174 | 338 | 797 | 1237 | 2385 |
| | PFM | 19 | 23 | 54 | 121 | 216 | 478 | 673 | 986 |
| | OFM | 25 | 39 | 78 | 164 | 321 | 743 | 1139 | 2074 |
| After filtering | ALL | 44 | 13 | 22 | 49 | 90 | 176 | 234 | 294 |
| | PFM | 19 | 11 | 20 | 45 | 81 | 157 | 203 | 234 |
| | OFM | 25 | 12 | 21 | 48 | 89 | 171 | 228 | 284 |

**Table S3** Annotation of OTUs with a number of reads greater than 1000 at the level of phylum, genus and species for the peach fruit moth (PFM) *Carposina sasakii* and the oriental fruit moth (OFM) *Grapholita molesta.*

| Phylum | Genus | Species | OTU code | Average depth | |
|---|---|---|---|---|---|
| | | | | PFM | OFM |
| Proteobacteria | *Wolbachia* | *Wolbachia* sp. | OTU461 | 3057.6 | 2.2 |
| | *Tatumella* | *Tatumella ptyseos* | OTU220 | 6.9 | 43.6 |
| | *Pseudomonas* | *Pseudomonas brenneri* | OTU2363 | 660.3 | 1673.4 |
| | | *Pseudomonas* sp. | OTU1066 | 266.9 | 735.0 |
| | | | OTU2143 | 12.7 | 32.3 |
| | *Pantoea* | *Pantoea* sp. | OTU226 | 158.4 | 283.9 |
| | | *Enterobacter ludwigii* | OTU1277 | 18.3 | 223.5 |
| | *Leuconostoc* | *Leuconostoc pseudomesenteroides* | OTU1039 | 31.6 | 20.8 |
| | *Lactobacillus* | *Lactobacillus iners* AB-1 | OTU2136 | 0.1 | 172.0 |
| | | *Lactobacillus* sp. | OTU1053 | 1.1 | 88.6 |
| | | | OTU1043 | 0.7 | 41.8 |
| | *Komagataeibacter* | *Komagataeibacter saccharivorans* | OTU499 | 42.3 | 59.0 |
| | *Gluconobacter* | *Gluconobacter cerinus* | OTU349 | 292.6 | 330.4 |
| | | *Gluconobacter albidus* | OTU1050 | 44.6 | 133.0 |
| | | *Gluconobacter frateurii* | OTU1074 | 6.7 | 153.8 |
| Firmicutes | *Frateuria* | *Frateuria aurantia* | OTU1067 | 10.2 | 74.9 |
| | *Dickeya* | *Dickeya* sp. | OTU350 | 12.4 | 101.0 |
| | *Acetobacter* | *Acetobacter* sp. | OTU1071 | 151.5 | 275.3 |
| | | | OTU1144 | 47.9 | 50.3 |

**Table S4** Details of bacterial genus shared among the peach fruit moth (PFM) *Carposina sasakii* and the oriental fruit moth (OFM) *Grapholita molesta.*

| Species | Phylum | Genus | PFM-19 samples | OFM-25 samples |
|---|---|---|---|---|
| PFM & OFM | Proteobacteria | *Pseudomonas* | 17870 | 61541 |
| | Proteobacteria | *Gluconobacter* | 6534 | 15429 |
| | Proteobacteria | *Pantoea* | 3363 | 13182 |
| | Proteobacteria | *Acetobacter* | 3789 | 8139 |
| | Proteobacteria | *Komagataeibacter* | 1044 | 2105 |
| PFM | Proteobacteria | *Wolbachia* | 58095 | 54 |
| OFM | Firmicutes | *Lactobacillus* | 124 | 9419 |
| | Proteobacteria | *Dickeya* | 236 | 2525 |

**Table S5** Alpha diversity of the gut bacterial microbiota for the peach fruit moth (PFM) *Carposina sasakii* and the oriental fruit moth (OFM) *Grapholita molesta.* Samples were numbered based on their population codes. See Table 1 for population codes. Pd: Phylogenetic diversity.

| Species | Sample | OTU | Sobs | Shannon | Simpson | Ace | Chao | Coverage | Shannon even | Simpson even | Pd |
|---|---|---|---|---|---|---|---|---|---|---|---|
| PFM | PDAE04 | 67 | 67 | 0.8610 | 0.5751 | 115.7458 | 104.4000 | 0.9931 | 0.2048 | 0.0260 | 12.8596 |
| | PDAE09 | 19 | 19 | 0.0595 | 0.9862 | 49.2312 | 30.0000 | 0.9978 | 0.0202 | 0.0534 | 3.5604 |
| | PDAE12 | 26 | 26 | 1.0913 | 0.4365 | 36.7373 | 37.2500 | 0.9980 | 0.3350 | 0.0881 | 3.2619 |
| | PDAE13 | 22 | 22 | 1.4862 | 0.2727 | 63.5276 | 34.0000 | 0.9982 | 0.4808 | 0.1667 | 3.0652 |
| | PDAE24 | 19 | 19 | 0.6874 | 0.6842 | 49.6524 | 28.3333 | 0.9984 | 0.2334 | 0.0769 | 2.4445 |
| | PGAE01 | 21 | 21 | 0.4196 | 0.8189 | 152.1000 | 47.0000 | 0.9974 | 0.1378 | 0.0582 | 3.9434 |
| | PGAE04 | 58 | 58 | 1.2631 | 0.3398 | 184.4177 | 102.0000 | 0.9933 | 0.3111 | 0.0507 | 7.6322 |
| | PGAE08 | 98 | 98 | 1.2646 | 0.4395 | 134.0886 | 137.3750 | 0.9927 | 0.2758 | 0.0232 | 14.7722 |
| | PGAE09 | 20 | 20 | 0.0548 | 0.9875 | 211.6869 | 140.0000 | 0.9968 | 0.0183 | 0.0506 | 3.7147 |
| | PKPR04 | 42 | 42 | 1.9008 | 0.2001 | 55.2214 | 50.6667 | 0.9974 | 0.5085 | 0.1190 | 5.7310 |
| | PKPR09 | 31 | 31 | 0.7312 | 0.7364 | 49.8736 | 42.0000 | 0.9976 | 0.2129 | 0.0438 | 3.9070 |
| | PKPR11 | 35 | 35 | 0.9665 | 0.5443 | 49.8941 | 46.0000 | 0.9976 | 0.2718 | 0.0525 | 4.0166 |
| | PKPR12 | 23 | 23 | 1.2628 | 0.3693 | 25.9591 | 24.0000 | 0.9992 | 0.4028 | 0.1177 | 2.9718 |
| | PKPR20 | 20 | 20 | 1.4874 | 0.3073 | 37.9776 | 25.0000 | 0.9988 | 0.4965 | 0.1627 | 3.3468 |
| | PLPR03 | 85 | 85 | 1.1500 | 0.5485 | 135.0146 | 126.3529 | 0.9923 | 0.2588 | 0.0214 | 9.7385 |
| | PLPR05 | 37 | 37 | 0.1707 | 0.9559 | 69.4000 | 60.3333 | 0.9957 | 0.0473 | 0.0283 | 5.3434 |
| | PLPR12 | 85 | 85 | 0.5854 | 0.8423 | 148.0391 | 114.5263 | 0.9931 | 0.1318 | 0.0140 | 12.5494 |
| | PLPR13 | 77 | 77 | 0.3636 | 0.9089 | 126.4089 | 108.5000 | 0.9927 | 0.0837 | 0.0143 | 14.1102 |
| | PLPR14 | 22 | 22 | 0.0900 | 0.9770 | 123.7885 | 57.0000 | 0.9970 | 0.0291 | 0.0465 | 3.6551 |
| OFM | OGAE01 | 63 | 63 | 2.0911 | 0.1622 | 95.8117 | 82.7143 | 0.9951 | 0.5047 | 0.0978 | 11.8754 |
| | OGAE02 | 74 | 74 | 1.1931 | 0.4536 | 95.5402 | 92.7500 | 0.9949 | 0.2772 | 0.0298 | 13.3562 |
| | OGAE03 | 135 | 135 | 2.3401 | 0.2010 | 181.0373 | 173.0769 | 0.9909 | 0.4771 | 0.0369 | 17.2739 |
| | OGAE06 | 73 | 73 | 1.0875 | 0.4821 | 102.9030 | 104.0000 | 0.9937 | 0.2535 | 0.0284 | 14.0081 |
| | OGAE09 | 63 | 63 | 1.3570 | 0.3695 | 95.4193 | 83.6471 | 0.9945 | 0.3275 | 0.0430 | 13.5850 |
| | OGAE11 | 29 | 29 | 1.0108 | 0.4556 | 48.5309 | 42.0000 | 0.9972 | 0.3002 | 0.0757 | 9.6853 |
| | OKPR11 | 43 | 43 | 1.7949 | 0.2349 | 78.1497 | 58.0000 | 0.9970 | 0.4772 | 0.0990 | 4.8230 |

| | | | | | | | | | | | |
|---|---|---|---|---|---|---|---|---|---|---|---|
| | OKPR12 | 57 | 57 | 2.3167 | 0.1647 | 132.3137 | 84.1429 | 0.9959 | 0.5730 | 0.1065 | 6.7532 |
| | OKPR13 | 53 | 53 | 2.1872 | 0.1561 | 81.6995 | 80.1429 | 0.9959 | 0.5509 | 0.1209 | 5.7518 |
| | OKPR19 | 34 | 34 | 1.7050 | 0.3260 | 40.5584 | 39.2500 | 0.9986 | 0.4835 | 0.0902 | 3.2704 |
| | OKPR24 | 13 | 13 | 0.5088 | 0.7438 | 17.7521 | 18.0000 | 0.9990 | 0.1983 | 0.1034 | 2.0355 |
| | OLPR01 | 71 | 71 | 1.6891 | 0.3112 | 91.7598 | 96.3000 | 0.9953 | 0.3962 | 0.0453 | 7.0046 |
| | OLPR09 | 118 | 118 | 2.2352 | 0.2075 | 145.3332 | 136.0968 | 0.9931 | 0.4685 | 0.0408 | 23.3562 |
| | OLPR12 | 137 | 137 | 2.1775 | 0.3353 | 157.7050 | 157.2174 | 0.9937 | 0.4426 | 0.0218 | 29.2981 |
| | OLPR13 | 90 | 90 | 1.7076 | 0.3159 | 119.8552 | 108.9000 | 0.9943 | 0.3795 | 0.0352 | 11.9615 |
| | OLPR14 | 144 | 144 | 2.9239 | 0.1527 | 165.7283 | 165.3684 | 0.9941 | 0.5883 | 0.0455 | 17.2300 |
| | OLPR15 | 64 | 64 | 1.1208 | 0.4820 | 124.9471 | 95.5000 | 0.9943 | 0.2695 | 0.0324 | 9.8241 |
| | OLPR17 | 101 | 101 | 1.6291 | 0.3663 | 142.0996 | 142.6250 | 0.9925 | 0.3530 | 0.0270 | 30.2145 |
| | OSPH06 | 55 | 55 | 0.5784 | 0.7678 | 141.9927 | 89.3636 | 0.9943 | 0.1443 | 0.0237 | 13.9065 |
| | OSPH09 | 42 | 42 | 0.9550 | 0.5035 | 57.9773 | 55.1538 | 0.9961 | 0.2555 | 0.0473 | 6.2602 |
| | OYPH07 | 52 | 52 | 0.9181 | 0.5207 | 69.1125 | 64.7500 | 0.9963 | 0.2324 | 0.0369 | 11.6263 |
| | OYPH12 | 40 | 40 | 0.8682 | 0.5265 | 55.4784 | 48.0769 | 0.9970 | 0.2354 | 0.0475 | 7.2079 |
| | OYPH14 | 56 | 56 | 1.1905 | 0.4522 | 77.6034 | 79.3333 | 0.9957 | 0.2957 | 0.0395 | 7.2816 |
| | OYPH19 | 35 | 35 | 0.8376 | 0.5513 | 53.4626 | 46.6667 | 0.9970 | 0.2356 | 0.0518 | 6.2456 |
| | OYPH28 | 31 | 31 | 1.8953 | 0.2131 | 125.5497 | 61.3333 | 0.9972 | 0.5519 | 0.1514 | 5.2653 |

**Table S6** Statistical tests on differences of alpha diversity between pairs of species/populations of the peach fruit moth (PFM) *Carposina sasakii* and the oriental fruit moth (OFM) *Grapholita molesta*. See Table 1 for population codes. Pd: Phylogenetic diversity.

| Test method | Type of index | Index | PFM-OFM | PGAE-OGAE | PKPR-OKPR | PLPR-OLPR | PDAE-PGAE | PKPR-PLPR | OKPR-OLPR | OSPH-OYPH |
|---|---|---|---|---|---|---|---|---|---|---|
| Student's t test | Richness | Sobs | 0.0164 | 0.3336 | 0.2981 | 0.0389 | 0.3661 | 0.0553 | 0.0021 | 0.5451 |
| | | Ace | 0.7780 | 0.0311 | 0.2310 | 0.3755 | 0.0017 | 0.0007 | 0.0082 | 0.4813 |
| | | Chao | 0.1366 | 0.7230 | 0.2141 | 0.0725 | 0.0490 | 0.0069 | 0.0014 | 0.4069 |
| | Diversity | Shannon | 0.0004 | 0.0763 | 0.2866 | 0.0008 | 0.8259 | 0.0209 | 0.5613 | 0.3285 |
| | | Simpson | 0.0006 | 0.0713 | 0.4820 | 5.789e-05 | 0.7811 | 0.0095 | 0.8868 | 0.2042 |
| | | pd | 0.0087 | 0.0431 | 0.5983 | 0.0686 | 0.4611 | 0.0395 | 0.0092 | 0.3906 |
| Welch's t test | Richness | Sobs | 0.0129 | 0.3485 | 0.3087 | 0.0401 | 0.4127 | 0.0782 | 0.0012 | 0.5476 |
| | | Ace | 0.7861 | 0.0267 | 0.2564 | 0.3963 | 0.0026 | 0.0028 | 0.0247 | 0.6724 |
| | | Chao | 0.1376 | 0.7254 | 0.2301 | 0.0847 | 0.0659 | 0.0147 | 0.0017 | 0.6025 |
| | Diversity | Shannon | 0.0004 | 0.0917 | 0.2923 | 0.0005 | 0.8300 | 0.0210 | 0.5802 | 0.2536 |
| | | Simpson | 0.0015 | 0.1517 | 0.4824 | 0.0008 | 0.7857 | 0.0101 | 0.9025 | 0.3732 |
| | | Pd | 0.0053 | 0.1064 | 0.6022 | 0.0478 | 0.4739 | 0.0637 | 0.0073 | 0.6230 |
| Wilcoxon rank-sum test | Richness | Sobs | 0.0142 | 0.2395 | 0.2963 | 0.0735 | 0.5386 | 0.1425 | 0.0058 | 0.5613 |
| | | Ace | 0.5696 | 0.0428 | 0.4034 | 0.5160 | 0.0200 | 0.0122 | 0.0230 | 0.5613 |
| | | Chao | 0.1207 | 0.5940 | 0.4034 | 0.1439 | 0.0662 | 0.0122 | 0.0058 | 0.5613 |
| | Diversity | Shannon | 0.0021 | 0.2410 | 0.2963 | 0.0094 | 0.9025 | 0.0367 | 1.0000 | 0.5613 |
| | | Simpson | 0.0036 | 0.3374 | 0.4034 | 0.0058 | 0.7133 | 0.0216 | 0.7453 | 0.5613 |
| | | Pd | 0.0060 | 0.1658 | 0.6761 | 0.1044 | 0.1113 | 0.0947 | 0.0058 | 0.5613 |

**Table S7** Significance analysis (Bray-Curtis, permutation = 999) of grouping factor for the peach fruit moth (PFM) *Carposina sasakii* and the oriental fruit moth (OFM) *Grapholita molesta* using PERMANOVA. See Table 1 for population codes.

| Characteristics | Sums of sequence | MeanSqs | F Model | R2 | P value |
|---|---|---|---|---|---|
| PFM&OFM | 3.6281 | 3.6281 | 15.2041 | 0.2658 | 0.0010 |
| PGAE/OGAE | 1.2500 | 1.2500 | 9.9348 | 0.5539 | 0.0040 |
| PLPR/OLPR | 1.8540 | 1.8540 | 9.3421 | 0.4830 | 0.0050 |
| PKPR/OKPR | 0.7008 | 0.7008 | 3.0381 | 0.2752 | 0.0350 |
| PLPR/PKPR | 0.4265 | 0.4265 | 3.9910 | 0.3328 | 0.0320 |
| PDAE/PGAE | 0.3010 | 0.3010 | 1.6607 | 0.1918 | 0.2330 |
| OLPR/OKPR | 0.5575 | 0.5575 | 1.8738 | 0.1578 | 0.0980 |
| PLPR/PKPR/PGAE/PDAE | 1.3176 | 0.4392 | 3.1022 | 0.3829 | 0.0150 |
| OLPR/OKPR/OGAE/OYPH | 1.6502 | 0.5501 | 2.5397 | 0.2862 | 0.0090 |

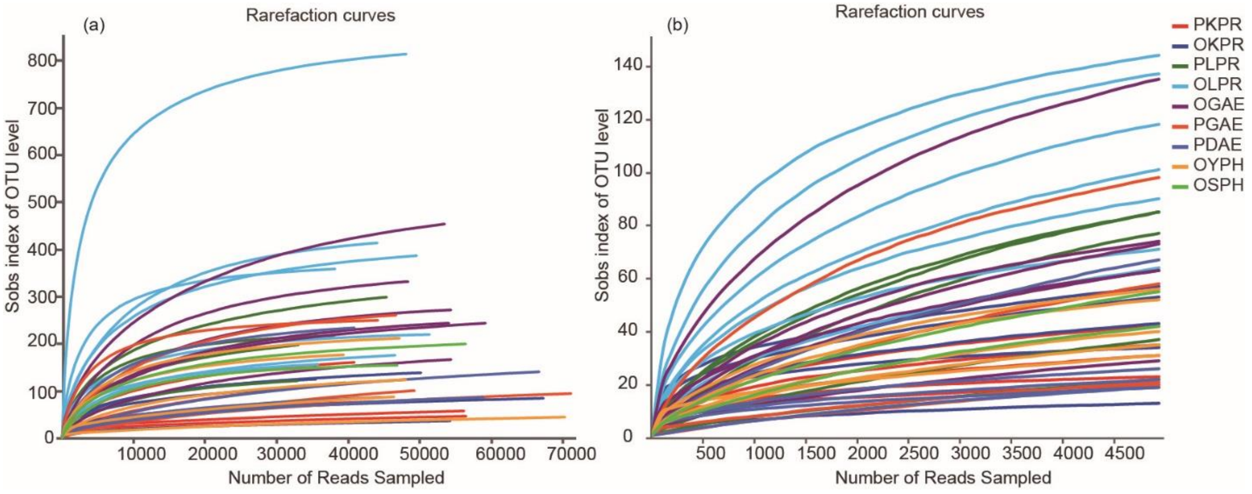


**Fig. S1** Rarefaction curves depicted from original sequencing data sets and randomly subsampled data sets with the same number of 16S rRNA sequences determined from the peach fruit moth (PFM), *Carposina sasakii*, and the oriental fruit moth (OFM) *Grapholita molesta*. (a) Rarefaction curves of original sequencing samples. (b) Rarefaction curves of subsampled sequences. See Table 1 for population code.

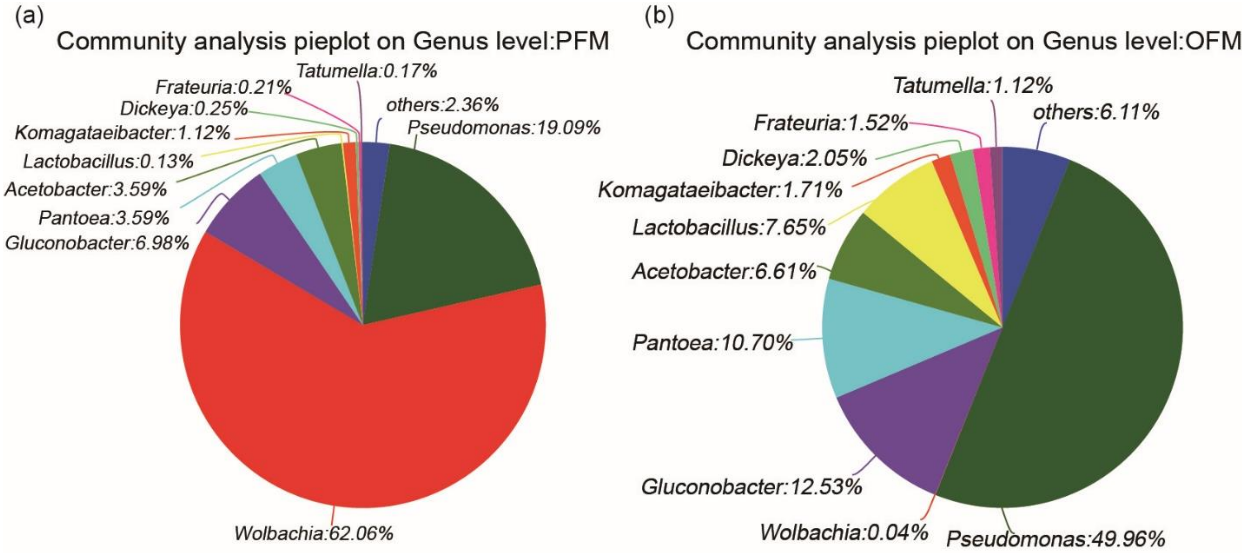


**Fig. S2** Community composition bacterial microbiota for the peach fruit moth (PFM), *Carposina sasakii*, and the oriental fruit moth (OFM) *Grapholita molesta*. (a)The composition of microbiome at the genus level in PFM. (b)The composition of microbiome at the genus level in OFM.

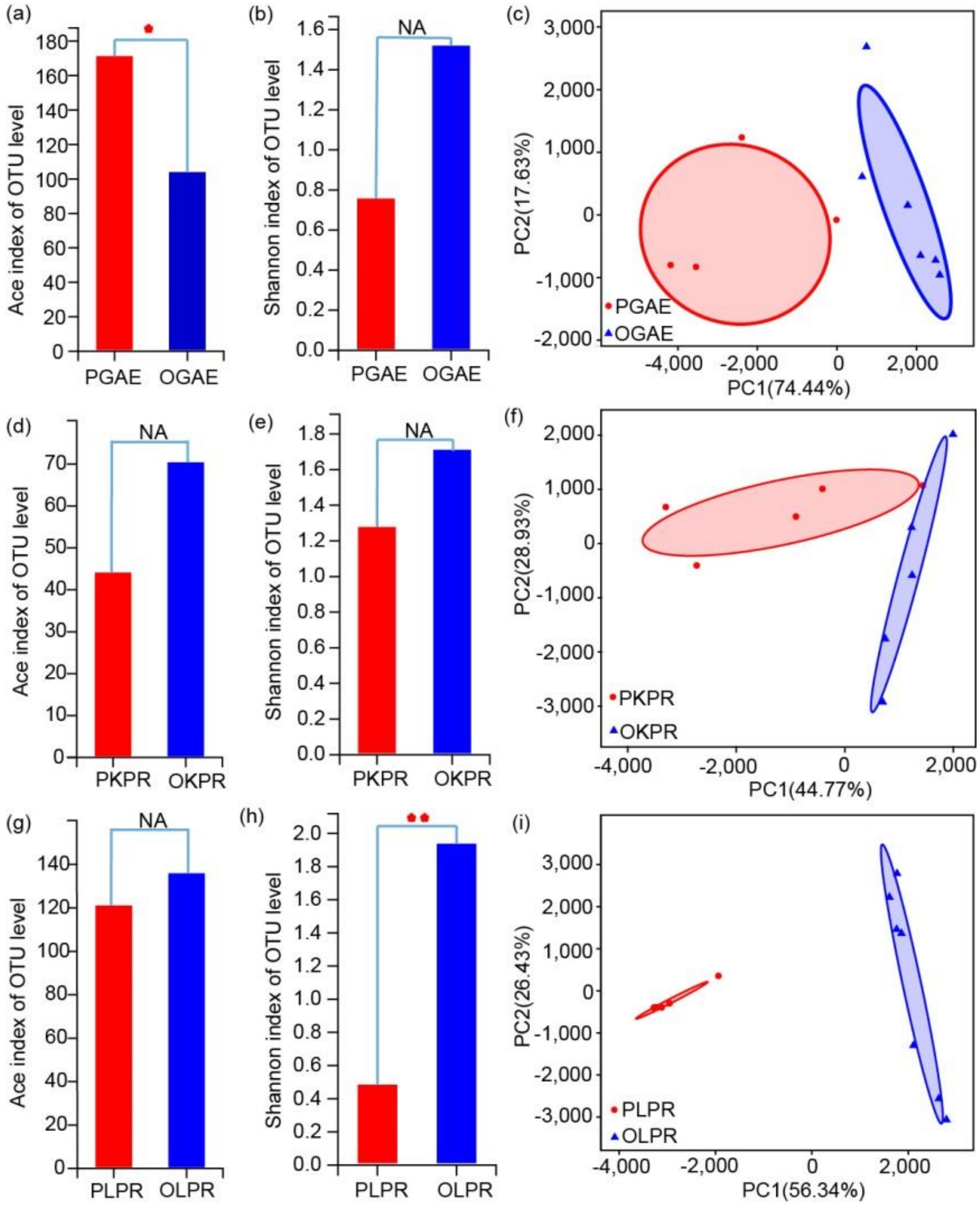


**Fig. S3** Difference in gut bacterial microbiota between the peach fruit moth (PFM), *Carposina sasakii*, and the oriental fruit moth (OFM) *Grapholita molesta* collected from the same fruit in an orchard. (a and b) Alpha diversity of microbiome between two species from the same host plant (apple) and orchard. A Wilcoxon rank-sum test compared microbiome richness based on the ace index and community diversity Shannon index between PFM (PGAE) and OFM (OGAE). (c) Beta diversity of the microbiome between the two species (PGAE and OGAE) analysed by PCA at the genus level. (d and e) Alpha diversity of PFM (PKPR) and OFM (OKPR) from the same pear orchard, and (f) associated PCA analysis of beta diversity. (g and h) Alpha diversity of the PFM (PLPR) and OFM (OLPR) microbiome in another pear orchard, and (i) associated PCA analysis of beta diversity. ($p > 0.05$ is marked as NA,$0.01 < P \leq 0.05$ is marked as *, $0.001 < p \leq 0.01$ is marked as **, and $P \leq 0.001$ is marked as ***).

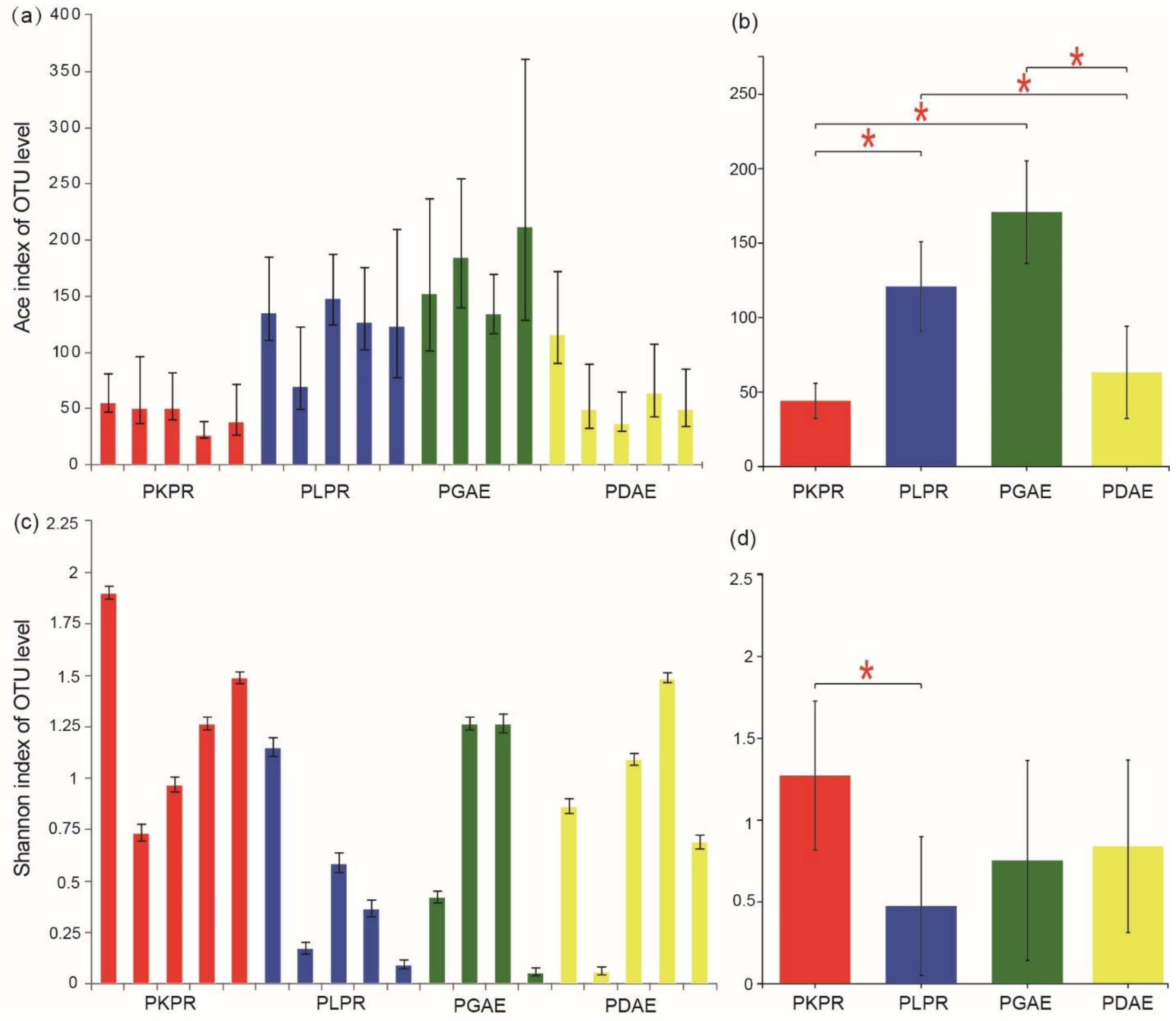


**Fig. S4** Alpha diversity of the microbiome of the peach fruit moth (PFM), *Carposina sasakii,* from four orchards. (a and c) At the OTU level, the richness (Ace) and diversity (Shannon) of individuals are plotted as histograms. (b and d) The differences between groups in richness and diversity were compared by Wilcoxon rank-sum tests (*$p < 0.05$). Error bars indicate standard deviations.

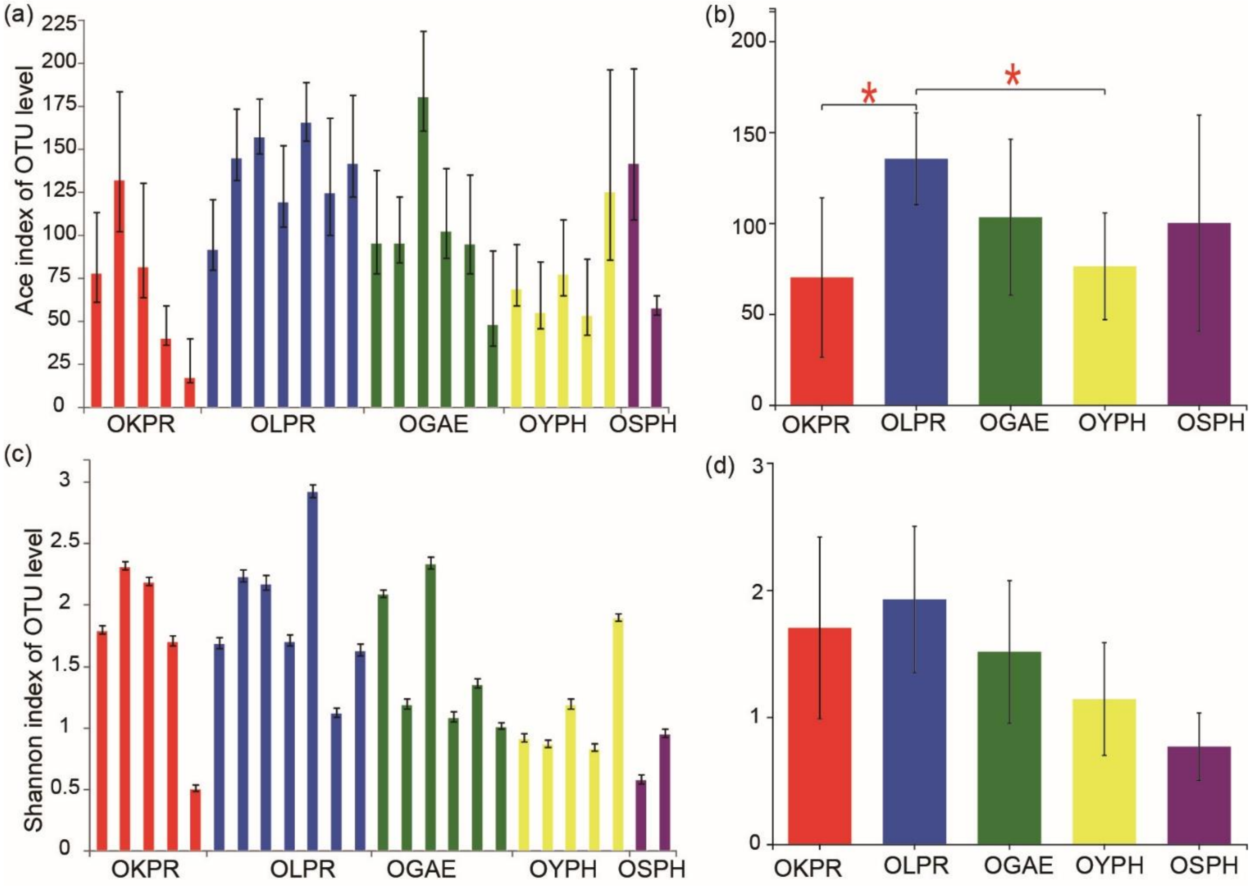


**Fig. S5** Alpha diversity of the microbiome of the oriental fruit moth (OFM), *Grapholita molesta,* from five orchards. (a and c) At the OTU level, the richness (Ace) and diversity (Shannon) of individuals are plotted as histograms. (b and d) The differences between groups in richness and diversity were compared by Wilcoxon rank-sum tests (*$p < 0.05$). Error bars indicate standard deviations.

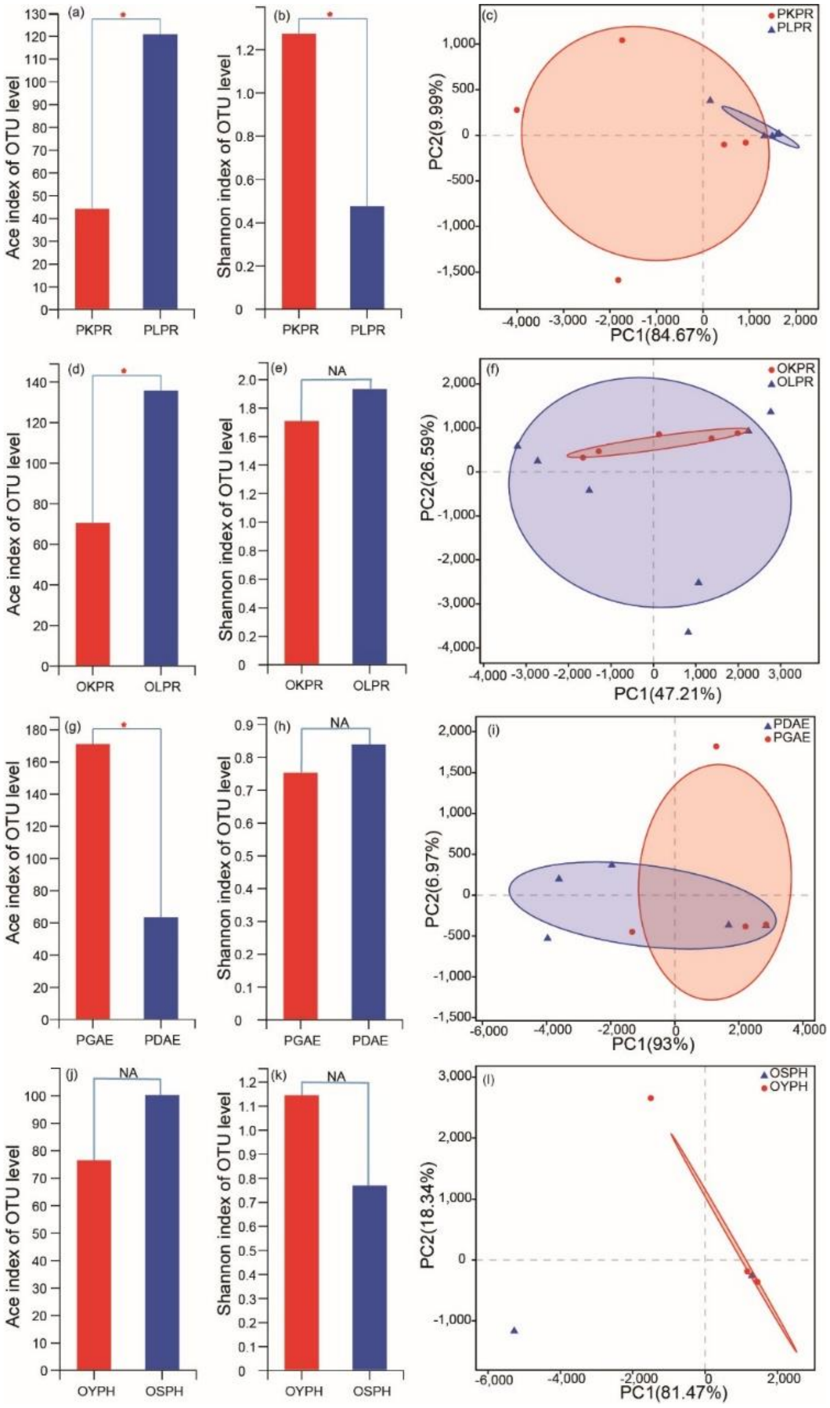


**Fig. S6** Difference of gut bacterial microbiota between orchards with the same host plant in the peach fruit moth (PFM), *Carposina sasakii*, and the oriental fruit moth (OFM) *Grapholita molesta*. Two pairs of orchards with the same host were compared for each species for alpha and beta diversity. (a and b) Wilcoxon rank-sum test of microbiome richness (Ace index) and community diversity (Shannon index) of the *Carposina sasakii* microbiome in two pear orchards (PLPR and PKPR). (c) Beta diversity of the microbiome between PLPR and PKPR estimated by PCA analysis at the genus level. (d and e) Alpha diversity of the *Grapholita molesta* microbiome in two pear orchards (OLPR and OKPR), with (f) providing the PCA analysis of beta diversity. (g and h) Alpha diversity analysis of the *Carposina sasakii* microbiome from two apple orchards (PGAE and PDAE) along with the PCA analysis of beta diversity (i). (j and k) Alpha diversity analysis of the *Grapholita molesta* microbiome in two apple orchards (OYPH and OSPH) along with the PCA analysis of beta diversity (l). ($P>0.05$ is marked as NA, $0.01 < P \leq 0.05$ is marked as *, $0.001 < P \leq 0.01$ is marked as **, and $P \leq 0.001$ is marked as ***).
